# GraPPI: A Self-Supervised Graph Encoder for Transferable Protein–Protein Interaction Modeling

**DOI:** 10.64898/2026.09.28.755140

**Authors:** Zhiyuan Song, Ziyu Shi, Gary Sun, Surendra Negi, Hugues Fausther-Bovendo, Haiqing Zhao

## Abstract

Protein–Protein Interactions (PPIs) are fundamental to cellular processes and represent major targets for therapeutic development. Recent advances in machine learning for protein modeling, particularly protein language models, have enabled powerful representations of protein sequences and monomeric structures. However, compared with the increasingly generalizable representations available for individual proteins, transferable representations explicitly designed to capture protein-complex and interface contexts remain less developed. Here, we developed **GraPPI**, a unified graph-based deep learning framework for learning transferable representations of protein–protein complexes. At its core, PPIencoder represents protein complexes as heterogeneous residual graphs and uses self-supervised masked-edge prediction to learn interaction-aware representations that integrate sequence, three-dimensional geometry, and physicochemical properties. We then evaluated the transferability of these pretrained representations across complementary PPI tasks spanning binding-mode classification, binding-affinity prediction, and mutation-associated affinity prediction. GraPPI accurately distinguished plausible binding interfaces from perturbed or decoy binding interfaces, achieving AUROC values above 0.92 on an independent benchmark. For protein–protein binding-affinity prediction, GraPPI achieved Pearson correlation coefficients of 0.690 and 0.684 on the S90 and S79 benchmarks, respectively, with performance competitive with or exceeding that of recently reported affinity-prediction methods. GraPPI further transferred effectively to mutation-associated binding-affinity prediction, achieving a Pearson correlation of 0.84 and a mean absolute error of 1.20 kcal/mol on an independent SKEMPI benchmark. Together, these results establish GraPPI as a generalizable framework for protein-complex representation learning, extending pretrained protein representations from individual proteins toward the interaction context in which protein function is realized.

## Introduction

Recent advances in machine learning (ML) have transformed protein science by enabling large-scale learning of biological representations from sequence and structural data. Protein language models (pLMs), such as ESM^1^ and ProtTrans^2^ are trained on large protein sequence databases and learn representations that capture evolutionary, structural, and functional information directly from amino-acid sequences^3,4^. In parallel, deep-learning approaches such as AlphaFold^5^ have dramatically advanced protein structure prediction. Together, these developments have established pretrained protein representations as powerful and transferable inputs for diverse protein-related prediction tasks.

Proteins, however, rarely function independently, and many biological processes depend on interactions between proteins and other biomolecules. Protein-Protein Interactions (PPIs) introduce a level of molecular organization that is not fully defined by either interacting protein alone: complex formation creates intermolecular contacts, binding-interface geometry, and physicochemical environments that emerge upon molecular association. Representing a protein complex therefore requires not only information about the individual protein sequences and structures, but also explicit description of the spatial and chemical relationships between interacting partners to inform the binding-interface geometry and physicochemical environments. Graph neural networks (GNNs)^6,7^ are particularly suited to this problem because protein complexes can be represented as residues or atoms connected through spatial and molecular relationships, allowing local geometry and interaction context to be incorporated directly into learned representations.

To address these challenges, a range of computational approaches have been developed for PPI modeling, progressing from classical machine-learning methods based on engineered features to deep-learning models that learn representations directly from protein sequences and structures^8^. Earlier binding-affinity methods, including ISLAND^9^ and PPI-Affinity^10^, combined engineered structural and physicochemical features with classical ML algorithms such as support vector regression (SVR)^11–13^. More recently, deep-learning approaches have enabled interaction patterns to be learned directly from protein-complex structures, with applications spanning protein-docking model assessment, binding-affinity prediction, and mutational effect prediction. Representative examples include energy-based GCN models such as EGCN^14^ for protein-docking assessment, ProAffinity-GNN^15^ and PPAP^16^ for binding-affinity prediction, and PPIformer^17^, GeoPPI^18^, Binding-ddG-Predictor^19^, and MT-TopLap^20^ for predicting mutation-associated changes in protein binding. Nevertheless, transferable representation learning at the level of protein complexes remains less developed. Pretrained pLMs provide broadly reusable representations of individual protein sequences but do not explicitly represent the geometry and physicochemical context created by protein-complex assembly. Conversely, many existing structure-based PPI models are optimized and evaluated for particular supervised prediction objectives. This leaves an important question: can interaction-aware representations be learned directly from protein complexes through self-supervised learning and subsequently transferred across distinct PPI prediction tasks?

Here, we developed **GraPPI**, a self-supervised graph-representation-learning framework for PPI modeling. At its core, PPIencoder represents protein complexes as heterogeneous residue graphs and integrates pretrained ESM-2 (650M) representations with three-dimensional geometry and physicochemical features. Through self-supervised masked-edge prediction, PPIencoder learns residue representations that encode binding-partner and interaction context, without requiring experimental affinity or downstream-task labels during pretraining. We evaluated the transferability of these representations across complementary tasks spanning binding-mode classification, absolute binding-affinity prediction, and mutation-associated affinity prediction. GraPPI achieved strong performance across independent benchmarks relative to sequence- and structure-based models, demonstrating that explicit protein-complex context provides information complementary to sequence-derived representations. Together, these results establish GraPPI as a transferable representation-learning framework for protein complexes and support self-supervised learning at the protein-complex level as a strategy for modeling diverse properties of PPIs.

## Results

### GraPPI Architecture and Self-Supervised Learning of Protein-Complex Representations

To construct a diverse dataset for self-supervised learning, we curated heterodimeric complexes from the Protein Data Bank (PDB)^21^, referred to as the dimer set, and complemented them with receptor–ligand-annotated complexes collected primarily from the PPB-Affinity^22^ and related affinity resources, referred to as the affinity set. After reviewing overlapping entries and annotations to ensure consistent definitions of protein–protein interfaces, the two sets were combined for GraPPI pretraining (Fig. 1A; see Methods for details). Each protein complex was represented as a heterogeneous residue graph, with the two interacting partners assigned distinct node types and spatial contacts represented as intra- or inter-chain edges. Node features integrate ESM-2 residue embeddings with physicochemical and structural-context descriptors, whereas edge attributes encode residue-pair geometry through relative orientations and inter-residue atomic distance distributions. We additionally constructed GraPPI-base, in which ESM-2 embeddings were replaced by amino-acid one-hot encodings while all other features were retained, allowing us to evaluate the contribution of pretrained sequence representations.

**Figure 1.**
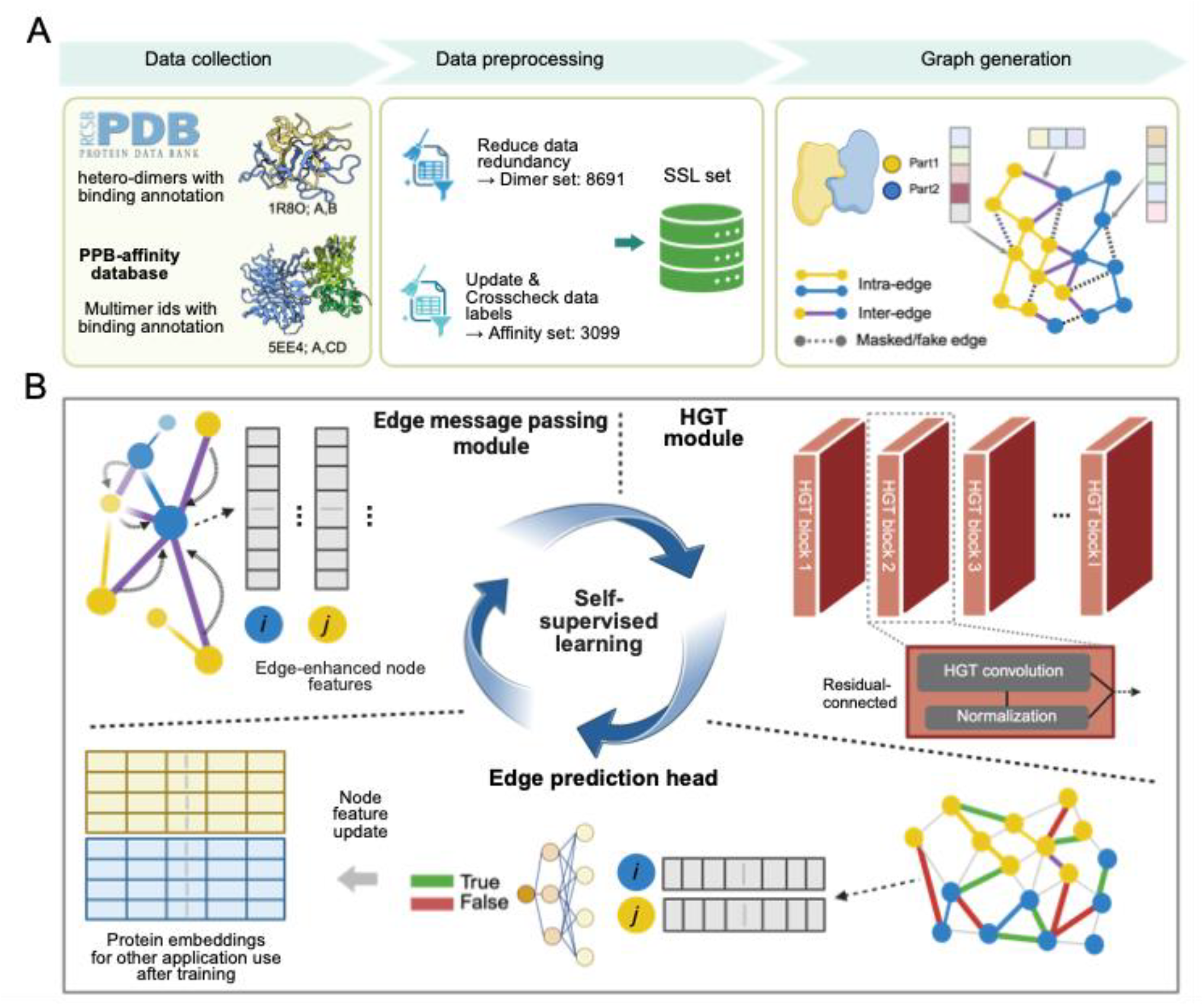
PPIencoder architecture and self-supervised pretraining. (A) Data curation and preprocessing workflow. Protein complexes were collected from PDB and PPB-Affinity database. PPB-Affinity labels and annotations were updated and cross-validated to complement the dimer complexes curated directly from PDB. (B) Model architecture of PPIencoder and self-supervised training. PPIencoder begins with an edge message-passing module that incorporates edge features into node feature updates (see Fig. S1 for details). The updated graph is then processed by a HGT module comprising multiple HGT blocks, each consisting of an HGT convolution layer residual-net connected by batch normalization layer. The resulting node embeddings are passed to an edge prediction head, which predicts the presence or absence of edges between node pairs, including masked or artificially generated edges. Model parameters are optimized through back-propagation using binary cross-entropy (BCE) loss.

PPIencoder, the representation-learning component of GraPPI, uses an edge-enhanced Heterogeneous Graph Transformer (HGT)^23^ to learn from the resulting protein-complex graphs (Fig. 1B; Methods). Because standard HGT does not directly incorporate edge attributes, we introduced an edge message-passing module that transforms geometric edge information through a gated multilayer perceptron^24^ (MLP), and then aggregates the resulting edge features into neighboring nodes in a relation-aware manner, followed by residual HGT blocks that learn interaction-context-dependent residue embeddings. PPIencoder was pretrained using self-supervised masked edge prediction (MEP)^25^ without experimental binding or affinity labels. During training, 25% of inter-chain edges and 15% of intra-chain edges connected to interface residues were masked, together with an equal number of generated negative edges, and the model was trained to recover edge relationships. The SSL dataset was divided into training and test sets at an 8:2 ratio.

We next evaluated the effects of model capacity and pretrained ESM-2 representations on self-supervised learning. Increasing HGT depth and embedding dimension generally reduced test loss, although the improvement was not monotonic with model size (Table S1). Both GraPPI and GraPPI-base showed decreasing training and test losses and increasing the area under ROC curve (AUROC) for edge-prediction during pretraining (Fig. S2). GraPPI-base exhibited a substantially higher initial test loss and slower convergence than GraPPI but ultimately reached similar test loss and AUROC. These results indicate that ESM-2 representations facilitate optimization, whereas the protein-complex graph itself provides sufficient structural and physicochemical information for learning the masked-edge prediction task.

Because performance on the self-supervised objective does not necessarily indicate that the learned representations are transferable, we next examined how training organizes the GraPPI embedding space and whether these representations support distinct PPI-related tasks, including binding-mode classification, binding-affinity prediction, and mutation-associated affinity prediction.

### Interaction-Associated Organization of GraPPI Representation

We first examined how self-supervised training shapes the residue-level representations learned by GraPPI. Because inter-chain contacts are explicitly represented in the input graph and targeted by the masked-edge prediction objective, we hypothesized that the learned representations would organize according to protein-interaction context. We first analyzed an immune signaling protein complex CD226**-**CD155 (PDB: 6ISC^26^), which was excluded from GraPPI pretraining, and projected its high-dimensional residue embeddings onto the first principal component (PC1) for visualization (Fig. 2A). GraPPI embeddings showed clear differences between interface and non-interface residues along PC1, whereas the original ESM-2 embeddings did not exhibit comparable separation (Fig. 2A and Fig. S3A). Importantly, randomly initialized GraPPI models (untrained) before training receiving the same graph inputs also failed to reproduce the interface-associated organization. Thus, although structural information defining inter-chain contacts is present in the input graph, its organization within the learned residue representations arises through self-supervised training rather than from the graph architecture or input features alone.

**Figure 2.**
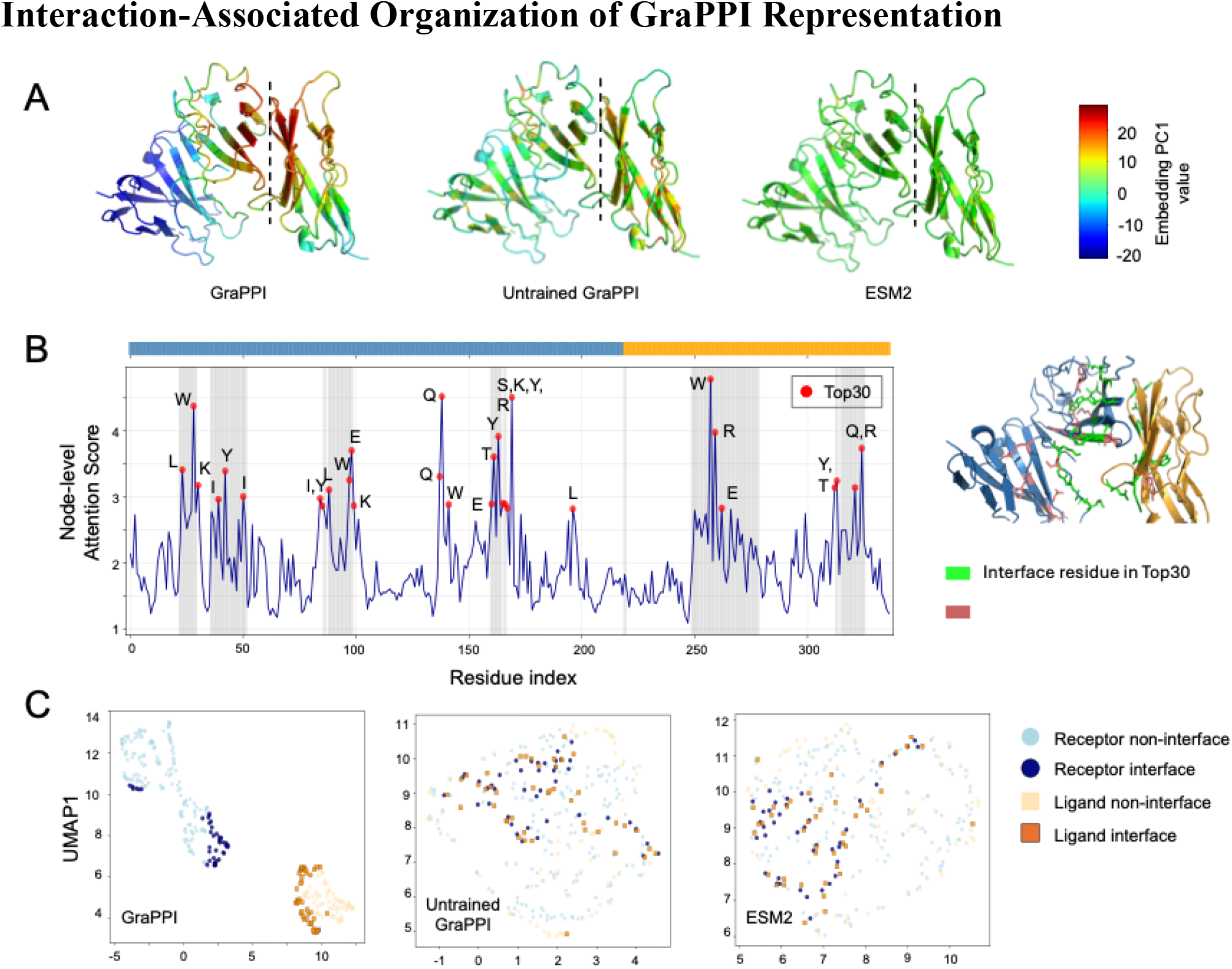
Visualization of residue-level embedding representations. (A) Residue embeddings from the immune signaling protein complex CD226-CD155 (PDB: 6ISC) were reduced to one dimension via PCA, and PC1 scores were mapped onto the protein structure to visualize the embedding patterns. The two proteins are separated by a dashed line. Results are shown for GraPPI, untrained GraPPI, and ESM-2 embeddings. (B) Residue-level attention scores extracted from the final HGT layer and mapped onto CD226-CD155 (blue/orange) sequences and structures. The top30 residues receiving the highest attention scores are labeled with red dots while native interface residues were marked in grey (left). (C) UMAP projection of residue embeddings from the CD226-CD155 complex, illustrating their organization in latent space. The two proteins are colored blue (CD226) and orange (CD155), with interface residues shown in a darker shade and non-interface residues in a lighter shade.

Because HGT uses attention mechanism to control information propagation across the protein-complex graph, we examined residue-level attention scores from the final HGT layer along the protein sequences (Fig. 2B). Among the 30 residues receiving the highest attention scores in CD226**-**CD155, 18 (60%) were interface residues, compared with an overall interface-residue frequency of 25.2% in the complex. This enrichment indicates that interface residues contribute disproportionately to information propagation within the trained GraPPI representation, consistent with the interaction-focused organization observed in the embedding analyses.

We further examined the organization of residue representations by projecting the embeddings into two dimensions using UMAP^27^ (Fig. 2C and Fig. S3B). GraPPI embeddings formed distinct clusters corresponding to the two interacting protein partners, with interface residues preferentially positioned toward regions connecting the two partner-specific distributions. In contrast, ESM-2 and untrained-model embeddings showed substantially less organization according to binding-partner and interface context. Together with the PCA analysis, these observations indicate that self-supervised training reorganizes residue representations to reflect both protein-partner identity and local interaction environment.

To test whether this observation generalized beyond the 6ISC example, we examined a set of 26 additional complexes: 24 randomly sampled from the test set (1%), none of which were used in GraPPI pretraining. For each complex, we compared PC1 distributions between interface and non-interface residues. Across these unseen complexes, trained GraPPI and GraPPI-base embeddings showed stronger statistical separation between interface and non-interface residues than ESM-2 embeddings or their corresponding untrained controls (Table S2). These results indicate that the interaction-associated organization observed for 6ISC is broadly reproduced across unseen protein complexes.

### GraPPI Representations Transfer to Binding-mode Classification

We first evaluated whether pretrained GraPPI representations could transfer to a downstream task involving recognition of plausible protein binding modes. This task is relevant to both experimentally determined and computationally predicted complexes, for which an observed interface may not necessarily represent a biologically relevant binding mode^28–30^. Because large-scale curated datasets of implausible protein–protein interfaces are not readily available, we constructed NegaPPI, a negative PPI dataset comprising three complementary sources: (1) low-confidence domain pairs derived from PrePPI-AF; (2) partner-swapped decoys generated by randomly exchanging binding partners among structurally dissimilar PDB dimers; and (3) interface-perturbed complexes in which interface residue identities and physicochemical features were altered while the backbone geometry was retained (Fig. 3A). Experimentally derived complexes were used as positive examples, and the independently constructed AbEpiTope^31,32^ swapped antibody–antigen dataset was additionally included for external evaluation. Dataset construction, filtering, and splitting procedures are described in the Methods.

**Figure 3.**
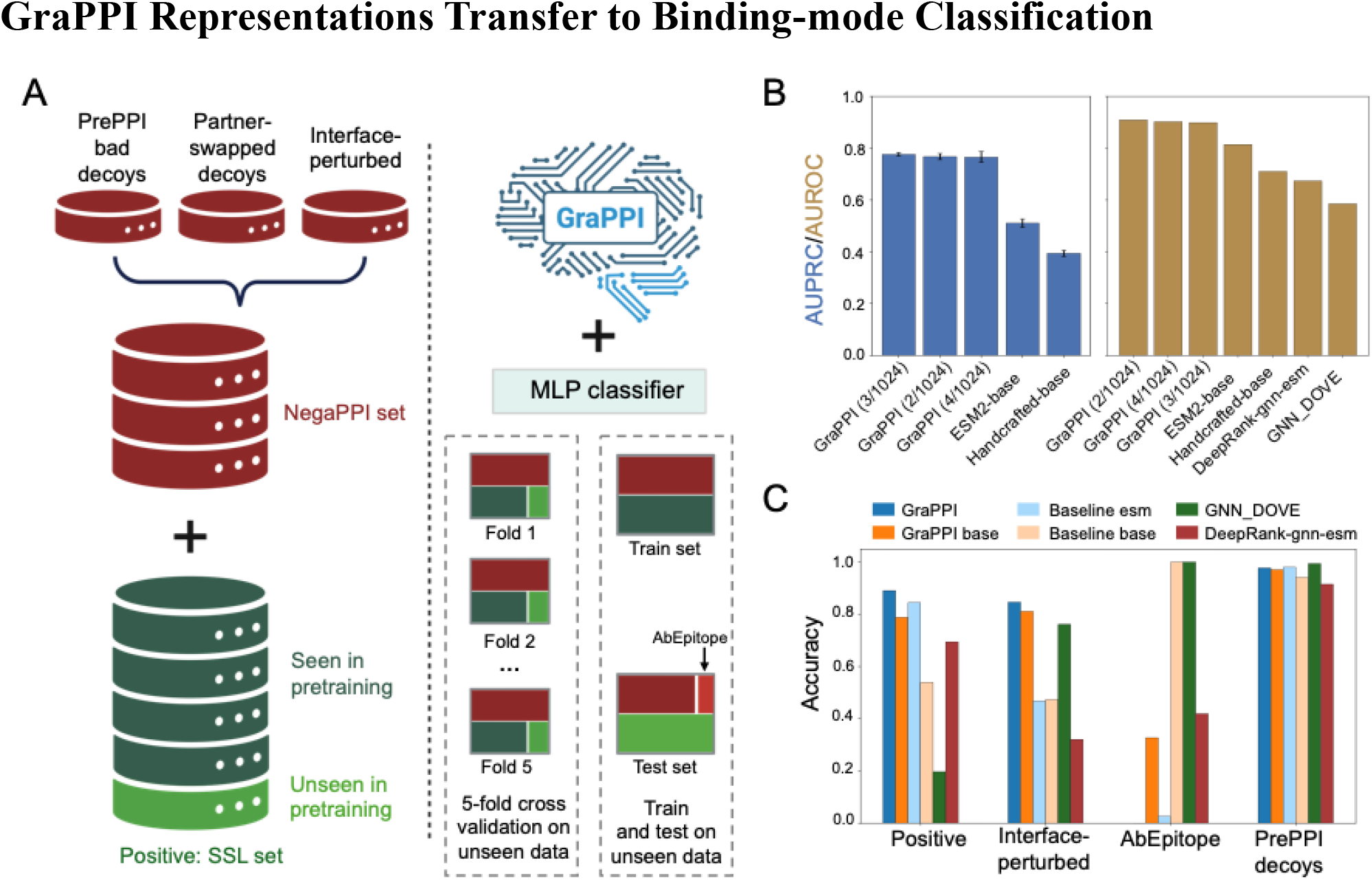
Dataset curation and model performance for the binding-mode classification task. (A) Construction of positive and negative dataset and data-splitting strategy for the binding-mode classification. (B) Classification performance of different GraPPI configurations and baseline models—GraPPI base and handcrafted-base—evaluated by 5-fold cross-validation and on the independent benchmark test set. Performance of GNN-DOVE and DeepRank-GNN-esm on the benchmark test set is included for comparison. (C) Classification accuracy of the selected GraPPI model across different benchmark subsets.

To transfer the pretrained residue representations to binding-mode classification, GraPPI embeddings were aggregated using Jumping Knowledge (JK-Net) aggregation^33^ and a cross-attention^34^ modules and classified using a two-layer MLP. We additionally evaluated conventional machine-learning classifiers such as SVM^35^ using the JK-Net outputs as input features (Table S3), and the constructed controls in which GraPPI representations were replaced by ESM-2 embeddings or the corresponding base features. In five-fold cross-validation, GraPPI models with 2, 3, and 4 HGT blocks and a hidden dimension of 1024 consistently achieved higher average precision (AUPRC) than the ESM-2 and base-feature controls (Fig. 3B), indicating that the pretrained complex representations provide information useful for distinguishing plausible from perturbed or decoy binding modes.

We next evaluated the models on the independent test set. As a control, we included two baseline models. The handcrafted-base model operates directly on handcrafted structural and physicochemical features, whereas the ESM2-base model uses ESM-2 representations without going through GraPPI pretraining. Both models employ the same downstream prediction framework, enabling direct comparison with GraPPI-derived embeddings (see Methods). The three selected GraPPI configurations achieved AUROC values above 0.92, with GraPPI (2/1024) achieving the highest test-set AUROC and therefore being selected for subsequent analysis (Fig. 3B). Across individual test subsets, GraPPI (2/1024) achieved accuracies of 0.89 for positive complexes and 0.85 for interface-perturbed complexes, outperforming GraPPI-base and the representation controls on these subsets (Fig. 3C). Together, these results show that the interaction-aware representations learned during self-supervised pretraining can be transferred to recognition of protein-complex binding modes not directly used as labels during pretraining.

One exception was the AbEpiTope subset, for which the base-feature model achieved higher classification accuracy than the GraPPI-based models. Examination of prediction-score distributions showed that the base-feature model produced a relatively narrow distribution concentrated at intermediate scores, whereas the other models exhibited more strongly separated score distributions (Fig. S4). Thus, the higher threshold-dependent accuracy of the base-feature model on this particular subset should be interpreted cautiously and does not correspond to stronger overall discrimination. Collectively, the cross-validation and independent-test results demonstrate that GraPPI representations transfer effectively from self-supervised protein-complex learning to binding-mode classification.

### GraPPI Representations Transfer to Quantitative Binding-Affinity Prediction

Following the binding-mode classification task, we next examined whether GraPPI representations capture quantitative properties of protein–protein interactions. We evaluated two complementary tasks: prediction of absolute protein–protein binding affinity and prediction of binding affinity for mutant complexes. Together, these tasks assess whether the pretrained representations can transfer from recognition of protein-complex binding modes to quantitative characterization of interaction strength and mutation-associated changes.

#### Protein–Protein Binding-Affinity Prediction

While binding-mode classification evaluates whether GraPPI can distinguish plausible from perturbed or decoy interfaces, it does not directly assess whether the learned representations encode quantitative properties of protein interactions. We therefore evaluated GraPPI for prediction of experimentally measured protein–protein binding free energy (ΔG). The affinity set was used for training and five-fold cross-validation. The S90 and S79 benchmarks were excluded from PPIencoder pretraining as well as from affinity-model training and were reserved solely for independent evaluation. GraPPI representations were transferred to the regression task using JK-Net aggregation and cross-attention, followed by a panel of regressors, including SVR, MLP, and other standard regression models; additional ablations are reported in Methods and Table S4.

Across five-fold cross-validation, GraPPI consistently performed better than GraPPI-base and the base-feature control, with GraPPI (6/1024) achieving the highest validation Pearson correlation (*R*_***P***_ = 0.661; Table 1). On the combined S90 and S79 independent test set, GraPPI (6/512) achieved *R*_***P***_ = 0.645 and the lowest MAE of 1.604 kcal/mol. The ESM2-base model achieved a comparable correlation (*R*_***P***_ = 0.615), whereas GraPPI produced a lower prediction error. The improvement over GraPPI-base further indicates that pretrained ESM-2 information and protein-complex structural context provide complementary information for quantitative affinity prediction.

**Table 1.** Model performance on the binding-affinity prediction task. Models were evaluated using 5-fold CV. *R*_*P*_ denotes the Pearson correlation coefficient. The Validation *R*_*P*_ column reports the mean *R*_*P*_ across the five validation folds, with values presented as mean ± standard deviation (std). The Test Rp/MAE column reports the *R*_*P*_ and means absolute error (MAE) obtained on the held-out test set using models trained on the remaining four folds. MAE values are reported in kcal/mol. GraPPI and baseline models here connect to SVR for the regression task.

| Model | No. HGT blocks | Feature dimension | Validation $R_P$ | Test $R_P$ (S90+S79) | Test MAE (S90+S79) |
| --- | --- | --- | --- | --- | --- |
| GraPPI | 6 | 1024 | $0.661 \pm 0.014$ | $0.615 \pm 0.021$ | $1.634 \pm 0.024$ |
| GraPPI | 6 | 512 | $0.658 \pm 0.017$ | $0.645 \pm 0.021$ | $1.604 \pm 0.019$ |
| GraPPI base | 2 | 1024 | $0.589 \pm 0.021$ | $0.583 \pm 0.019$ | $1.735 \pm 0.026$ |
| GraPPI base | 2 | 512 | $0.568 \pm 0.021$ | $0.549 \pm 0.020$ | $1.786 \pm 0.025$ |
| ESM2-base | None | 1280 | $0.600 \pm 0.012$ | $0.615 \pm 0.013$ | $1.663 \pm 0.010$ |
| Handcrafted-base | None | 25 | $0.406 \pm 0.028$ | $0.551 \pm 0.008$ | $1.836 \pm 0.011$ |

We next compared GraPPI (6/512) with previously published methods on the S90 and S79 benchmarks (Table 2) using the best test outcome among the 5 folds. GraPPI achieved *R*_***P***_ = 0.690 and MAE = 1.47 kcal/mol on S90 and *R*_***P***_ = 0.684 and MAE = 1.69 kcal/mol on S79. On the combined benchmark, GraPPI achieved *R*_***P***_ = 0.685 and MAE = 1.57 kcal/mol. These results place GraPPI among the competitive methods across both benchmarks and, importantly, demonstrate that representations learned through self-supervised protein-complex pretraining can be transferred to quantitative binding-affinity prediction.

**Table 2.**
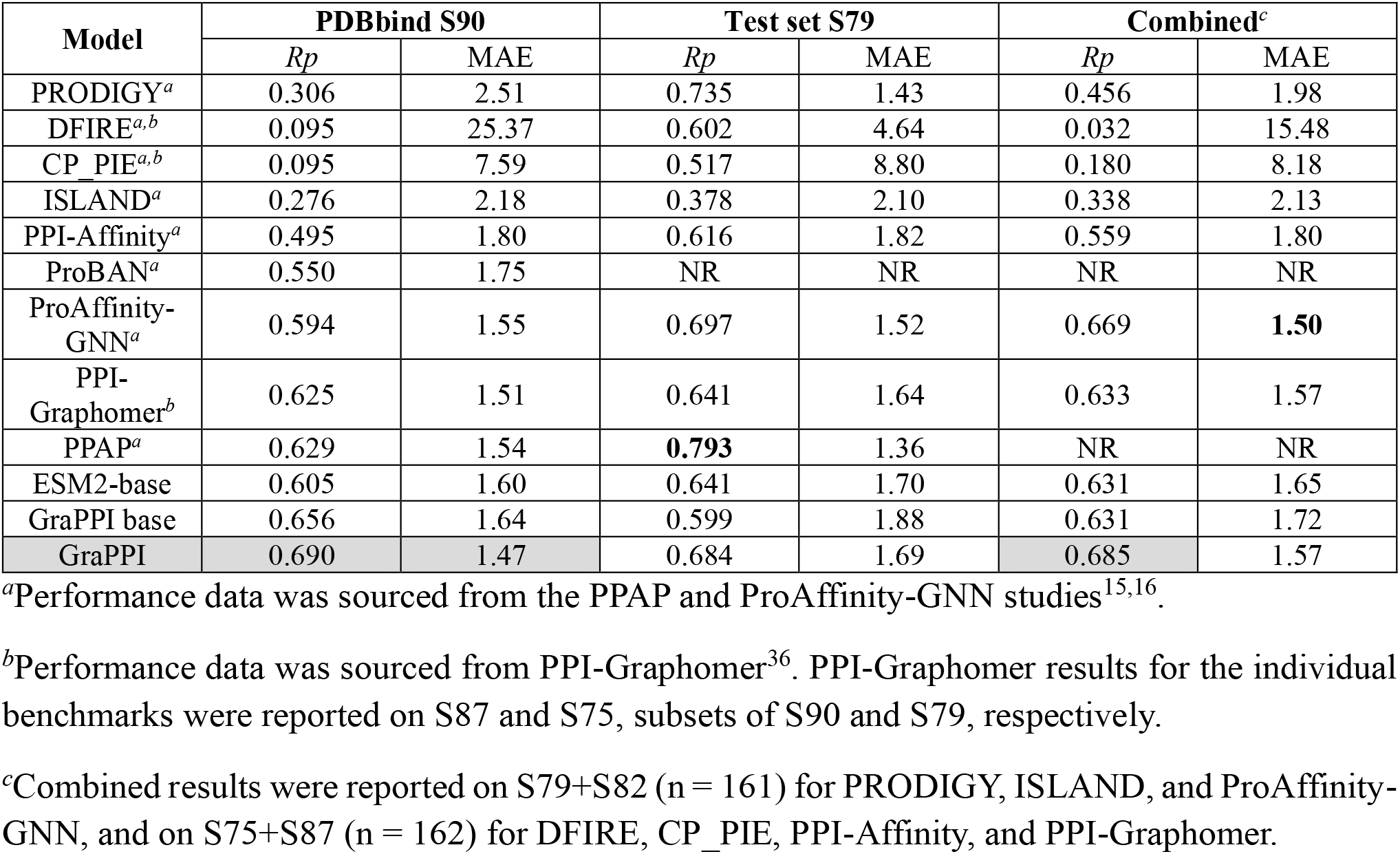
Comparison of GraPPI with published models for protein–protein binding-affinity prediction. Performance is reported using *Rp* and MAE (kcal/mol) on two widely used benchmark datasets, PDBbind S90 and S79. GraPPI was evaluated on the complete S90, S79 and their union (n = 169), whereas some published results were reported on subsets (see footnotes). “NR” indicates data not reported.

#### Mutation-Associated Binding-Affinity Prediction

We next asked whether GraPPI representations could transfer to the more localized perturbations introduced by interface mutations. The mutation subset of PPB-Affinity, containing 5,019 samples, was used for training and cross-validation, and an independent SKEMPI set containing 26 wild-type complexes and 140 corresponding mutants — none of which overlapped with the PPIencoder pretraining set — was used for testing. Mutant structures were generated using FoldX^37^. Using the same general transfer-learning framework, models were trained to predict the experimentally measured binding free energy (mutant ΔG) of mutant complexes (see Methods and Table S5).

During cross-validation, GraPPI achieved *R*_***P***_ = 0.859, compared with 0.826 for GraPPI-base and 0.824 for ESM-2 (Table 3). On the independent SKEMPI benchmark, GraPPI achieved *R*_***P***_ = 0.817 and MAE = 1.298 kcal/mol. ESM-2 produced a slightly higher correlation (*R*_***P***_ = 0.838) but a higher MAE (1.353 kcal/mol), whereas GraPPI-base showed lower performance by both metrics (*R*_***P***_ = 0.717; MAE = 1.413 kcal/mol). Thus, sequence-derived representations remain highly informative for mutation-associated affinity, while incorporating protein-complex geometry and physicochemical context improves quantitative prediction error and maintains strong correlation on an independent benchmark.

**Table 3.** Model performance on the mutational binding-affinity prediction task. The validation *R*_*P*_ column reports the mean *R*_*P*_ across the five validation folds, with values presented as mean ± std. The Test Rp/MAE column reports the *R*_*P*_ and MAE obtained on the held-out test set using models trained on the remaining four folds. MAE values are reported in kcal/mol

| Model | No. HGT blocks | Feature dimension | Validation $R_P$ | Test $R_P$ | Test MAE |
| --- | --- | --- | --- | --- | --- |
| GraPPI | 2 | 1024 | $0.859 \pm 0.011$ | $0.817 \pm 0.016$ | $1.298 \pm 0.060$ |
| GraPPI base | 2 | 256 | $0.826 \pm 0.014$ | $0.717 \pm 0.023$ | $1.413 \pm 0.021$ |
| ESM2-base | None | 1280 | $0.824 \pm 0.015$ | $0.838 \pm 0.007$ | $1.353 \pm 0.042$ |
| Handcrafted-base | None | 25 | $0.687 \pm 0.024$ | $0.218 \pm 0.023$ | $2.197 \pm 0.040$ |

Comparison with previously published approaches further showed that GraPPI remained competitive on the SKEMPI benchmark, achieving *R*_***P***_ = 0.84 and MAE = 1.198 kcal/mol in the corresponding benchmark evaluation (Table 4) using the best performed fold for test set. Notably, several compared methods were developed specifically for mutation-effect prediction, whereas GraPPI uses a general protein-complex representation learned without affinity or mutation labels during pretraining. Together, the absolute and mutation-associated affinity results demonstrate that GraPPI representations transfer across distinct quantitative PPI prediction problems and complement sequence-derived representations with explicit protein-complex context.

**Table 4.** Comparison of GraPPI with published models for mutation-associated protein–protein binding-affinity prediction. Performance was evaluated on the widely used benchmark dataset, SKEMPI test set. Superscripts indicate metric data sourced from ^*a*^PPAP, ^*b*^PPI-Affinity, or ^*c*^Pro-Affinity-GNN. “NR” means data not reported in the previous publications.

| Model | SKEMPI test set |  | Mutation-specific model |
| --- | --- | --- | --- |
| | $R_P$ | MAE | |
| PPIformer | $0.46^a$ | NR | Yes |
| GeoPPI | $0.52^a$ | NR | Yes |
| Binding-ddg-predictor | $0.73^a$ | NR | Yes |
| SSIPE | $0.61^a$ | NR | Yes |
| MT-TopLap | $0.88^a$ | NR | Yes |
| FoldX | 0.38 | 3.422 | Yes |
| PPI-Affinity | $0.78^b$ | $1.4^b$ | No |
| Pro-Affinity-GNN | $0.73^c$ | $1.964^c$ | No |
| ESM2-base | 0.85 | 1.270 | No |
| PPAP | $0.80^a$ | 1.440 | No |
| GraPPI base | 0.72 | 1.388 | No |
| GraPPI | 0.84 | 1.198 | No |

**Table 5.** Summary of heterogeneous graph components.

| Heterogeneous Graph Element | Description |
| --- | --- |
| Node type $\mathcal{T}$ | receptor, ligand |
| Relation types $\mathcal{R}$ | (receptor → receptor), (receptor → ligand), (ligand → ligand), (ligand → receptor) |
| Node feature dimension $\mathbf{x}_v^{(\tau)}$ | 1285, 25 (base version) |
| Edge feature dimension $\mathbf{e}_{uv}^{(r)}$ | 8 |

**Table 6.** Summary of supported JK-Net mode details.

| JK-Net mode | Aggregation function: $\text{Agg}(\cdot)$ | Output Dimension: $d_{\text{jk-net}}$ |
| --- | --- | --- |
| Mean (Actual mode used in GraPPI) | $\frac{1}{L+1} \sum_l \mathbf{H}^l$ | $d_{\text{hidden}}$ |
| Concat (Optional) | $[\mathbf{H}^{(0)} \parallel \dots \parallel \mathbf{H}^{(L)}]$ | $d_{\text{hidden}} \times (L+1)$ |

## Discussion

In this study, we developed GraPPI as a self-supervised representation-learning framework designed specifically for protein–protein complexes. Whereas pretrained protein language models provide powerful representations of individual protein sequences, protein interactions additionally depend on the three-dimensional geometry and physicochemical environment formed upon complex assembly. GraPPI addresses this complementary level of representation by modeling protein complexes as heterogeneous residue graphs and explicitly incorporating inter-residue geometry and interaction context during self-supervised learning. The resulting representations showed interaction-associated organization and transferred across distinct downstream tasks spanning binding-mode classification, absolute binding-affinity prediction, and mutation-associated affinity prediction.

The performance across these tasks provides insight into the information captured by GraPPI representations. Representations learned by PPIencoder through self-supervised masked-edge prediction encode the binding-partner identity and interface context more effectively than either ESM-2 embeddings or an untrained PPIencoder. More importantly, these representations transferred to prediction targets that were not supplied as labels during pretraining. GraPPI achieved strong discrimination of plausible versus perturbed or decoy binding modes and competitive performance on quantitative binding-affinity benchmarks. The comparison with ESM-2 is particularly informative: GraPPI improved both correlation and prediction error for absolute binding affinity, whereas ESM-2 achieved a slightly higher correlation for mutant-complex affinity while GraPPI produced a lower prediction error. Rather than indicating universal superiority of either representation, these results suggest that sequence-derived information and explicit protein-complex context capture complementary aspects of protein interactions. This complementarity may be particularly useful for developing generalizable models that connect protein sequence, complex structure, and interaction energetics.

Several limitations arise from the structural data used for representation learning. Although GraPPI was pretrained on more than 11,000 protein complexes, experimentally determined structures represent a limited and non-random sample of protein-interaction space. PDB structures are influenced by biological interest and experimental tractability, potentially overrepresenting certain proteins, interaction classes, and interface architectures. Individual structures may also contain truncated constructs, engineered mutations, fusion partners, or conformations stabilized under non-physiological conditions. Automated curation reduces but cannot eliminate these sources of bias and uncertainty. The continuing expansion of experimentally determined structures, together with carefully filtered high-confidence predicted complexes, should substantially broaden the structural space available for future protein-complex representation learning.

Uncertainty in downstream supervision presents an additional challenge. For binding-mode classification, large collections of experimentally validated incorrect binding modes are not available, requiring us to construct complementary negative examples through partner swapping, low-confidence PrePPI-AF candidates, and interface-feature perturbation. These examples provide useful computational controls but cannot be considered experimentally established non-interactions in every case. Binding-affinity labels introduce a different source of uncertainty: structural models and affinity measurements may correspond to different constructs or experimental conditions, and measurements for the same interaction can vary among studies. During dataset curation, overlapping affinity measurements from independently curated resources showed a Pearson correlation of approximately 0.85, illustrating the substantial experimental and curation-related variability in available affinity data (Fig. S5). Such label uncertainty may place a practical ceiling on improvements obtained solely through model development. GraPPI representations themselves also remain only partially interpretable. Although embedding and attention analyses reveal interaction-associated organization, they do not establish which molecular determinants are responsible for individual predictions. More systematic attribution and mechanistic analyses will therefore be important for connecting learned representations to physical principles governing protein recognition and binding energetics.

GraPPI provides a general framework that can evolve as protein-complex data and modeling methods improve. Expanding pretraining to broader experimentally determined and carefully selected predicted complexes, incorporating higher-quality quantitative interaction measurements, and developing more interpretable representations should further increase its applicability. More broadly, our results support protein complexes as a distinct and useful level of representation learning: sequence-derived representations describe properties intrinsic to individual proteins, while complex-aware representations can additionally encode the geometric and physicochemical context that emerges upon molecular association. Integrating these complementary levels may provide a path toward more general models of protein interactions and their perturbation by sequence, structural, and environmental changes.

## Methods

### Data Curation

#### Construction of the Affinity Set

Protein complex datasets such as PDBbind^38^ and SKEMPI^39^ have been widely used for developing models for protein–protein interaction analysis. However, these datasets are typically curated for specific downstream tasks, such as binding affinity or mutational effect prediction, and therefore primarily include complexes with experimentally measured binding properties. As a result, they are relatively limited in size and diversity, potentially introducing sampling bias during model training.

To augment the dimer set with complexes containing more than two chains, we built on the recently published PPB-Affinity database, which unifies entries from ATLAS^40^, SAbDab^41^, PDBbind, and SKEMPI. We parsed PPB-Affinity into a wild-type subset and a mutant subset and standardized all affinity measurements to ΔG (kcal/mol). When ΔG was not provided directly, *K*_D_ was converted via ΔG = −*RT* ln*K*_D_ using the reported temperature, or 298.15 K when temperature was missing. We then re-checked each of the four upstream databases for recent releases as of March 2026. SAbDab provided substantial new entries. They were added when the antigen was a protein or peptide, and a valid affinity value was available; for complexes that overlapped with PPB-Affinity, conflicting affinity values were reconciled by averaging. Finally, we compared the resulting set to the dataset used in ProAffinity-GNN. Conflicting affinity values for the same entries were reconciled by averaging and we further incorporated any complexes present there but absent from our collection. The final table constitutes the affinity set comprising 3099 complexes.

#### Construction of the Heterodimer Set

To assemble a structurally diverse pool of protein–protein complexes, we first mined hetero-dimeric structures from the RCSB PDB. The RCSB Search API was queried for entries whose oligomeric state was annotated as “Hetero 2-mer”, whose polymer type was protein, and which were determined either by X-ray diffraction or cryo-electron microscopy at a resolution of 4.0 Å or better, or by solution NMR. For each hit, the experimental structure was downloaded in PDB format, falling back to mmCIF (and converting to PDB with Biopython^42^) when the legacy PDB format was not available. Only standard amino-acid residues were retained, eliminating heteroatom ligands whose residue names exceed three characters and would otherwise corrupt the fixed-width PDB columns. Protein chains were identified as chains containing at least one Cα atom; when more than two protein chains were present (i.e., the asymmetric unit contained several symmetry-related copies of the same biological dimer), the biological dimer was resolved by a three-stage fallback. The RCSB biological-assembly REST endpoint (/core/assembly/{pdb}/1) was queried first, and the curator-annotated assembly was accepted when it returned exactly two chains matching the deposited protein chains; otherwise the REMARK 350 BIOMT records of the downloaded PDB file were parsed as a fallback; and if neither source produced an unambiguous pair, the first two protein chains in the deposited coordinates were retained as a last resort, with the resulting dimer subsequently re-validated by the sequence-identity and residue-contact checks described below.

Because the RCSB annotation of “Hetero 2-mer” does not guarantee a biologically interacting pair, every candidate dimer was further validated by two criteria. First, pairwise sequence identity between the two assigned chains was required to be below 0.95 to ensure that the pair was genuinely hetero. Second, the two chains were required to form at least five inter-chain residue contacts, where a contact was defined as any heavy-atom pair within 6.5 Å (computed efficiently with a KD-tree). For structures with more than two protein chains, the chain pair maximizing the number of contacts while satisfying the hetero criterion was selected. Candidates failing either filter were flagged for manual inspection rather than silently kept. The mined and validated dimers were then merged with an in-house pre-collected dataset of 5,037 hetero-dimers^43^, yielding more than 20,000 candidate hetero-dimeric complexes.

To remove redundancy, the candidates were clustered directly at the complex level with Foldseek’s easy-multimercluster^44^, using a coverage threshold of 0.80 and a multimer TM-score threshold of 0.80 (exhaustive search enabled to retain small proteins). Clustering at the complex level — rather than at the level of individual chains or on sequence — ensured that both partners and their relative orientation contributed to the similarity measure, which we found more appropriate for protein–protein interaction data than sequence-only clustering. For each Foldseek cluster, the medoid (the member minimizing the average TM-distance to the rest of the cluster) was retained as the cluster representative. Pairwise TM-score and RMSD matrices used both for medoid selection and for downstream analyses were precomputed with USalign^45^. The resulting set of cluster representatives constitutes the dimer set of 8691 samples.

#### Self-Supervised Learning Set

The self-supervised pre-training set, hereafter the SSL set, was defined as the union of the dimer set and the affinity set after removing complexes that appear in both. Crucially, every complex belonging to any downstream test set —the S90 and S79 ΔG benchmarks, the SKEMPI mutational test set, and the binding-mode classification test set (see below) — was removed from the SSL set prior to pre-training, resulting in 11585 samples. This guarantees that the encoder never observes a test complex, in either pre-training or fine-tuning.

#### Binding-mode Classification Dataset–NegaPPI set

We constructed a benchmark dataset to evaluate the ability of GraPPI embeddings to discriminate between plausible and implausible protein-protein interfaces. The binding-mode classification task requires positive samples representing plausible protein-protein interfaces and negative samples representing implausible interfaces. Positive samples were obtained from the SSL dataset. Because no large-scale, high-quality curated dataset of implausible protein-protein interfaces is available, we constructed the NegaPPI set with three complementary sources (Fig. 3A).

Source 1: low-confidence domain pairs from PrePPI-AF. We mined the human PrePPI-AF^46^ database for domain–domain pairs assigned a low PrePPI score (score between 5 and 10), which the PrePPI model itself deems unlikely to bind. To guard against false negatives, any pair was discarded whenever the two parent proteins were experimentally reported to interact in any form — even when the specific domain pair had no experimental evidence. The retained domain pairs were then assembled into 3D complexes using USalign, aligning to the PrePPI template structures. To exclude trivially non-binding complexes or ones with extensively steric clashes, the assembled complexes were required to display at least five inter-chain residue contacts (heavy-atom distance < 6.5 Å) and no more than five clashing residue pairs (heavy-atom distance < 2 Å); chain and complex sizes were further capped at 1,600 residues to keep the inputs computationally tractable. This procedure yielded up to 15,000 high-quality PrePPI-derived negatives, while 3104 of them were randomly sampled for GraPPI binding-mode classification task.

Source 2: partner-swapped decoys. Starting from the affinity set, we generated mismatched complexes by swapping ligand and receptor between different complexes. To avoid trivially similar negatives, the wild-type complexes were first clustered with hierarchical clustering using a combined TM-score/RMSD distance by USalign, average linkage, and a distance threshold of 0.65. One swap was generated per cluster, yielding 3,075 swapped complexes, generated with AlphaFold2-multimer.^47,48^

Source 3: interface-perturbed complexes. In contrast to the swap and PrePPI negatives, the interface-perturbed negatives were not realized as new three-dimensional structures. For each real complex, we identified the binding-interface residues on each partner, randomly selected five interface residues per partner (ten per complex), and mutated each to a different amino acid sampled uniformly from the remaining nineteen. The corresponding graph node features — the amino-acid one-hot encoding and the physicochemical descriptors — were updated accordingly, while the graph topology, the edge index and the edge attributes of the wild-type complex were retained. This graph-level perturbation mimics aggressive interface mutagenesis at the level the encoder actually consumes, presenting it with a complex whose interface chemistry has been heavily disrupted while the underlying backbone geometry remains intact, and avoids the computational cost and the structure-prediction artefacts that would be incurred by re-folding every mutated complex. Of the 8691 heterodimers, 136 lacked interface residues sufficient to support the perturbation procedure, leaving 8555 interface-perturbed complexes; of these, 3,089 were randomly sampled for the GraPPI binding-mode classification task to match the similar sample size of other data sources.

#### External Benchmark—AbEpiTope

As a stringent external test set for binding-mode classification, we used the publicly available AbEpiTope^31,32^ collection, which contains 272 swapped antibody–antigen complexes derived from 17 original Ab–Ag complexes and generated with AlphaFold2-multimer.

#### Mutation Subset

For the mutational ΔG task, the mutation subset of PPB-Affinity served as the training and validation set with 5019 samples, and the SKEMPI test of 26 wild type complexes and 140 corresponding mutants split was used as the held-out benchmark^15^. All mutant complex structures were generated with FoldX.^37^

### Graph Representation

To preserve binding-partner identity within each complex, the two interacting proteins were designated as receptor and ligand, following conventional protein–protein docking terminology, and their residues were assigned to two distinct node types. This assignment defines four directed edge relations (receptor→receptor, ligand→ligand, receptor→ligand, and ligand→receptor), enabling the model to distinguish intramolecular from intermolecular residue interactions and to support relation-specific information propagation within and across the two binding partners. Each protein–protein complex is represented as a heterogeneous graph *G* = ({***V***^(*τ*)^}_*τ*∈*T*_, {*ε*^(*r*)^}_*r*∈*R*_), where *T* = {receptor, ligand} is the set of node type; *R* = {*τ*_*s*_, *ϕ, τ*_*t*_} is the set of directed relation types with source type *τ*_*s*_, target type *τ*_*t*_, and relation label *ϕ*.

The base version of GraPPI encodes protein amino acid types using the one-hot representation as partial model inputs for node feature, e.g. (1, 0, 0, …) for Ala. To incorporate evolutionary information, we employed ESM-2 embeddings as an important component of the residue node features. Specifically, protein sequence embeddings were generated using the pretrained ESM-2 model (ESM2-t33-650M-UR50D). Given a protein with sequence length of N, the model produces a residue-level embedding of dimension N×1280. Each row of this embedding corresponds to a single residue and is directly used as the core component of the node feature representation.

To incorporate binding-context and physicochemical information, additional residue-level features were appended to the ESM embeddings. First, the binding partner identity of each residue was encoded using a one-hot vector of size 2, indicating which binding component the residue belongs to. Second, residue charge was encoded using discrete values {-1, 0, 1}, for negatively charged, neutral, and positively charged residues. Third, residue hydrophobicity was encoded according to the hydrophobicity scale reported by Rose *et al*..^49^ Finally, based on the chemical structure of each amino acid, a binary indicator was included to denote the presence or absence of a ring structure. In total, each node is represented by a feature vector of dimension of 1285, consisting of 1280-dimensional ESM embedding and 5 additional physicochemical and contextual features. These node features serve as the input to the GraPPI model.

Edges are constructed based on spatial proximity: an undirected edge is added between two residues if any atomic pair is closer than an 8 Å distance cutoff. Each node *v* ∈ ***V*** is associated with a feature vector 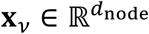, and each edge (*u, v*) ∈ *ε* is associated with an edge attribute vector 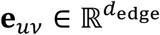 and a relation type *r* ∈ *R*.

Although the two inter-edge relations correspond to the same underlying pairwise contacts in opposite directions, they are retained as distinct directed relations to support direction-specific message passing between the two binding partners. Each edge is further associated with geometric attributes describing the spatial relationship between the connected residues. These attributes include (i) the unit vector pointing from the center of mass of residue *μ* to that of residue *v*, and (ii) a normalized histogram of interatomic distances between all atom pairs of the two residues. The distance histogram uses a bin width of 3 Å and covers the range from 0 to 15 Å.

### Heterogeneous Graph Transformer–based PPIencoder

#### Edge-Enhanced Message Passing

All node features of a graph are first projected into a unified hidden space (Fig. S1)

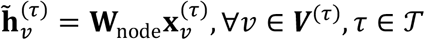

Where 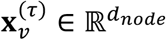 is the input node feature and 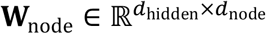 is the shared projection matrix (Same weights for all *τ* ∈ *T*; receptor and ligand nodes share the same input dimension *d*_node_).

Prior to graph transformer layers, GraPPI performs an edge message passing step to explicitly encode edge attributes. For each edge (*u,v*) with relation type r and the corresponding attributes 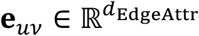, an edge message 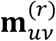 is computed using a relation-specific multilayer perceptron (MLP) with a bottle neck architecture with a gated modulation of source node features by edge attributes

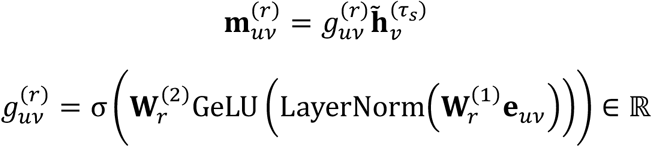

Where *σ* is the sigmoid function; the gate 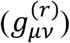 is a scalar that modulates the source node feature, 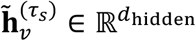 conditioned on edge attributes of relation *r*; 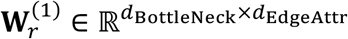 and 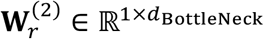 are linear projection matrices.

For each target node 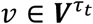, messages from all incoming relation types are aggregated

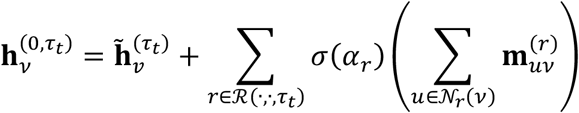

Where *N*_*r*_(*v*) is the set of source neighbors of *v* under relation *r* = (*τ*_*s*_, *ϕ, τ*_*t*_); *R*(·,·, *τ*_*t*_) = {*r* ∈ *R* | *r* = (*τ*_*s*_, *ϕ, τ*_*t*_)} selects only relations whose target type is *τ*_*t*_; *α*_*r*_ is a learnable scalar per relation type, initialized small.

#### Heterogeneous Graph Transformer

The edge-enhanced node embeddings, 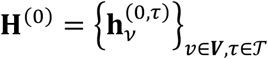 are further processed by a stack of *l* HGT blocks. Each block receives the full heterogeneous node embedding collection and the graph structure (edge indices per relation type) and returns updated node embeddings. Internally, HGTConv applies type-specific and relation-specific Q/K/V projections and relation-aware multi-head attention, guided by the metadata supplied at construction. The residual addition, shared LayerNorm, and edge dropout are applied externally in the encoder’s forward pass:

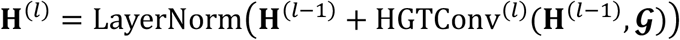

Where *G* denotes the heterogeneous graph structure, i.e. metadata. LayerNorm uses shared weights but is applied independently to each node type’s embedding matrix.

### Self-Supervised Learning of PPIEncoder

#### Masked Edge Prediction

Self-supervised pre-training was performed using a masked edge prediction (MEP) objective. A fixed proportion (25%) of inter-edges were randomly masked from the input graph. To further increase the task difficulty and encourage the model to learn interface-surrounding geometric and biochemical features, we additionally randomly masked 15% of intra-edges that connect at least one interface node, i.e., a node participating in at least one inter-edge.

To formulate a balanced binary classification task for edge prediction, we also generated an equal number of fake edges corresponding to the masked edges, enabling the model to predict both positive (true) and negative (false) edge existence.

For each candidate edge (*u, v*), a feature vector is constructed by concatenating the node embeddings **h**_*u*_ and **h**_*v*_, their absolute difference, and their element-wise product. This representation is passed through a 2-layer edge prediction head to produce a binary classification prediction logit. The following equation formulation is applicable to cases where **h**_*u*_ and **h**_*v*_ correspond to either the same node type or different node types.

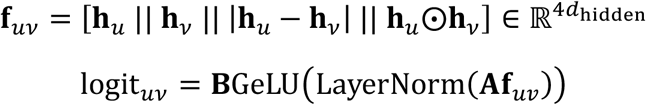

Where 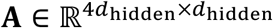 and 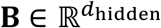 are learnable projection matrices, ⨀ denotes element-wise multiplication, || refers to concatenation operation. Dropout is applied after each layer during training to improve robustness.

#### Inter/Intra Combined Loss

SSL loss is a combination of *L*_intra_ and *L*_intra_. Each term is computed using binary cross-entropy (BCE) on the corresponding logits over masked positive edges and randomly sampled negative edges. *β* here is a tunable parameter.

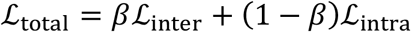

### Attention Analysis of PPIEncoder

The encoder uses a scaled dot-product attention scheme adapted from HGT. For each directed edge (*j→i*) of relation type *r* in HGT layer of block *l*, the unnormalized attention score for head *h* is:

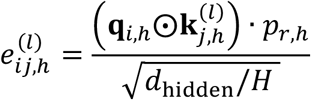

Where ⨀ denotes element-wise multiplication and *H* refers to the number of attention heads. 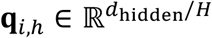 if the query vector of destination node *i* at block *l* for head h. 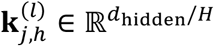 refers to the key vector source node *i* at block *l* for head h. *p*_*r,h*_ is one scalar per head per relation type, shared by all edges of that relation; initialized to 1 and optimized during training.

The raw scores are normalized across all incoming neighbors of *i* using SoftMax:

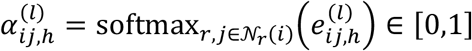

With the normalization constraint:

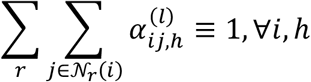

For every directed edge (*j→i*) of any relation, the multi-head attention 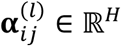 is first collapsed to a scalar by averaging over heads:

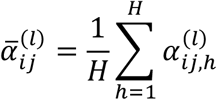

Then, we can obtain an attention score for a given node by aggregating incoming and outgoing attention scores:

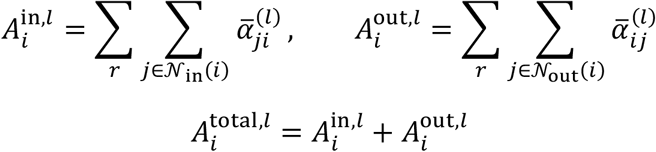

If we plug in the normalization constraint,

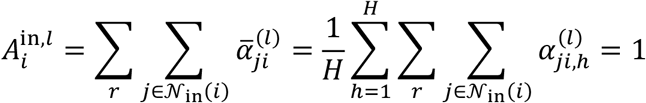

The informative quantity is therefore 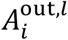.

### Application of the Pretrained PPIencoder

#### Jumping Knowledge Networks (JK-Net)

We applied JK-Net to aggregate node embeddings from different HGT blocks in order to: (1) capture multi-scale structural representations across network layers; (2) avoid imposing prior assumptions regarding the most relevant representation scale for a given downstream task; and (3) reduce the variance of the learned representations. The equations and table below illustrate how JK-Net was applied in this study:

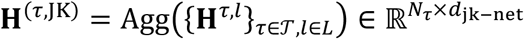

Where *N*_*τ*_ denotes the number of type *τ* nodes.

#### Pool Head for Dimension Reduction

A pool head is applied to the output from the JK-net:

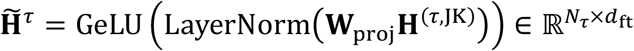

Where 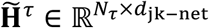 are the JK-aggregated node embeddings. *d*_ft_ = min(512, *d*_jk-net_/2)

#### Cross-Attention Aggregation

To further aggregate residue-level node embeddings into a single protein-complex representation with reduced dimensionality, we employed a cross-attention mechanism.

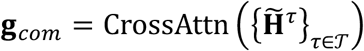

Specifically, we defined a set of learnable query vectors **Q**^(*τ*)^ ∈ ℝ^*Q*×*dft*^, which attend over the node embeddings of each node type through a two-stage multi-head attention framework. Query vectors **Q**^(*τ*)^ are learned during the specific downstream tasks.

In stage 1, we used an independent query-to-node attention to process pretrained protein complex embeddings:

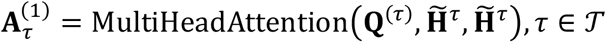

In stage 2, we used a residual network for for 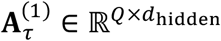 each node type:

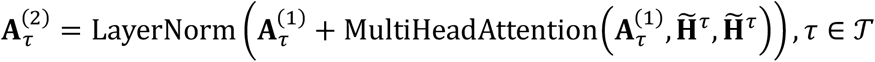

In stage 3, we combined the output from stage 2 to get the protein-level embedding for a given protein complex:

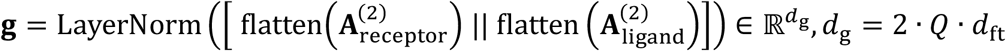

#### Task-specific MLP Prediction Heads

For all three downstream tasks, we designed an *K* layer MLP framework while individual parameters are in use. A certain input vector **g**_in_ can be further mapped to a scalar with a halving bottleneck. The forms of **g**_in_ and the MLP output are task-specific:

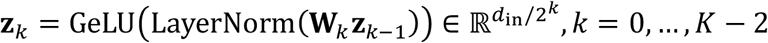

When *k=0*, ***z***_−1_ ≡ **g**_in_; *k=K-2*, the output dimension is forced to be 1. The details of different tasks are summarized in Table 7.

**Table 7.** Summary of model configurations for downstream GraPPI applications.

| Applications | Input $\rightarrow \mathbf{g}_{\text{in}}$ | Dimension: $d_{\text{in}}$ | Loss function | Validation metric |
| --- | --- | --- | --- | --- |
| Binding-mode classification | $\mathbf{g}$ | $d_g$ | BCE | AUROC |
| $\Delta G$ | $\mathbf{g}$ | $d_g$ | logcosh | MAE/Pearson $r$ |
| Mutational $\Delta G$ | $[\mathbf{g}_{\text{mut}} - \mathbf{g}_{\text{wt}} \parallel \mathbf{g}_{\text{mut}} \parallel \mathbf{g}_{\text{wt}}]$ | $3d_g$ | logcosh | MAE/Pearson $r$ |

#### Downstream Baseline Models

To assess the contribution of pretrained protein-complex representations, we also constructed two baseline models. The handcrafted-base model utilizes handcrafted protein-complex descriptors, including structural and physicochemical features of dimension of 25 per residue, as inputs to either conventional ML algorithms or shallow MLP. This model serves as a naïve baseline representing traditional feature-engineering approaches. In contrast, the ESM2-base baseline model replaces GraPPI embeddings with residue representations derived directly from ESM-2 of dimension of 1280 per residue and applies the same downstream prediction pipeline. This design enables evaluation of the benefits of protein language model representations without graph-based interaction learning. Together, these baselines allow us to distinguish the contributions of handcrafted features, sequence-derived embeddings, and GraPPI-learned protein-complex representations.

#### Evaluation Using Diverse Machine Learning Models

In addition to MLPs, we employed various ML algorithms, including support vector machine/regressor (SVM/SVR), Gradient Boosting, Random Forest, and Decision Tree models, across all application tasks. The ML models take the inputs from JK-net, 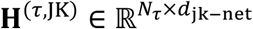, and obtain a representing vector via a mean pool for a given protein complex.

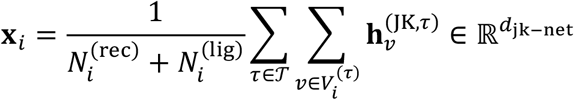

The embedding **x**_i_ is directly used as input for the ΔG prediction and binding-mode classification models. For mutational ΔG prediction, a concatenated representation of the mutant and wild-type embeddings, 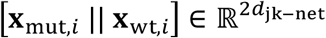, is used as the model input. Table 8 summarizes all tuned hyperparameters used in the machine learning models; parameters not listed were kept at their default values. In all, these complementary models enabled a comprehensive evaluation of the generalizability and effectiveness of GraPPI embeddings across diverse downstream learning architectures.

**Table 8.** Summary of hyperparameters for downstream machine-learning models.

| | Binding-mode classification | $\Delta G$ | Mutational $\Delta G$ |
| --- | --- | --- | --- |
| **Random forest** |  |  |  |
| n estimators | 500 | 300 | 300 |
| Max depth | 15 | 10 | 12 |
| Min samples leaf | 5 | 10 | 5 |
| Max features | sqrt | sqrt | sqrt |
| **Gradient boosting** |  |  |  |
| n estimators | 500 | 200 | 300 |
| Max depth | 6 | 4 | 5 |
| Learning rate | 0.1 | 0.05 | 0.1 |
| **Decision tree** |  |  |  |
| Max depth | 15 | 8 | 10 |
| Min samples leaf | 20 | 15 | 10 |
| **SVM/SVR** |  |  |  |
| SVM C | 1 | 10 | 10 |
| SVM epsilon | Default | 0.1 | 0.1 |

#### Training Protocol and Hyperparameters

All experiments were carried out on a single workstation equipped with NVIDIA GeForce RTX 5090 GPU and CUDA 12.8. The software stack is mainly PyTorch 2.7 (CUDA 12.8 build) with PyTorch Geometric and Scikit-learn. Mixed-precision training is enabled by default on CUDA devices via torch.amp.autocast with bfloat16, which reduces GPU memory. Model and SSL pre-training runs additionally use gradient checkpointing on the HGT stack. AdamW^50^ optimizer is employed for the training with decoupled weight decay. The learning rate is annealed by cosine annealing with warm restarts^51^. The minimum learning rate is set to be 10^−6^. Cosine warm-restart period *T*_*0*_ and multiplier *T*_mult_ are configurable and set to be 10 and 2, respectively. To accommodate larger gradient magnitudes early in training while tightening as gradients stabilize, the clipping threshold is updated adaptively after each step, *c*_*t*+1_ = ma**x**(*c*_min_, *α*||**g**_t_||_2_). *c*_min_ =5.0 and *α*=1.25. The clipped gradient **g**_*t*_ ← **g**_*t*_ · min(1, *c*_*t*_/|| **g**_*t*_||_2_). Training is halted when the validation metric for a given task fails to improve for a specified number of consecutive epochs (patience). The monitored metric varies by task. For SSL training, the edge-prediction test loss is monitored. For downstream MLP tasks, validation AUROC is used for binding-mode classification, while Pearson correlation (*R*_*P*_) is used for both ΔG and mutational ΔG prediction tasks.

## Data availability

The datasets generated and curated in this study have been deposited in Zenodo. The GraPPI-SSL dataset, comprising curated protein–protein complexes used for self-supervised pretraining, is available at https://zenodo.org/records/22946815. The NegaPPI dataset, comprising curated negative and decoy protein–protein complexes for model development and benchmarking, is available at https://zenodo.org/records/22968504.

## Code availability

The GraPPI source code, including codes for data preprocessing, model training, evaluation, and downstream analyses, is publicly available at https://github.com/zhaolabutmb/GraPPI.

## Acknowledgements

We thank Dr. Bernard M. Pettitt for helpful discussions. H.Z. acknowledges the generous support of the Welch Foundation (H-2308-20260402) and the UT System Rising STARs Award.

## Author contributions

Z.S and H.Z. designed research; Z.S., Zi.S., G.S. performed research; H.F.-B. contributed to data curation, Z.S., Zi.S., S.N., and H.Z. analyzed data; Z.S. and H.Z. wrote the manuscript; all authors revised the manuscript.

