## Supplemental Figures for "GraPPI: A Self-Supervised Graph Encoder for Transferable Protein–Protein Interaction Modeling"

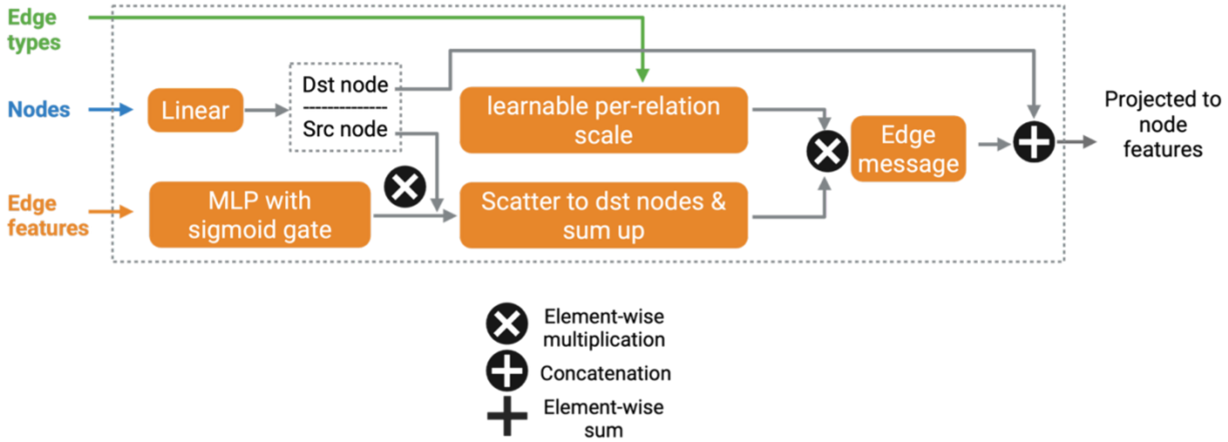


**Figure S1. Architecture of the Edge-Enhance module.** This module uses edge features to enhance node features before the graph is processed by the Heterogeneous Graph Transformer (HGT).

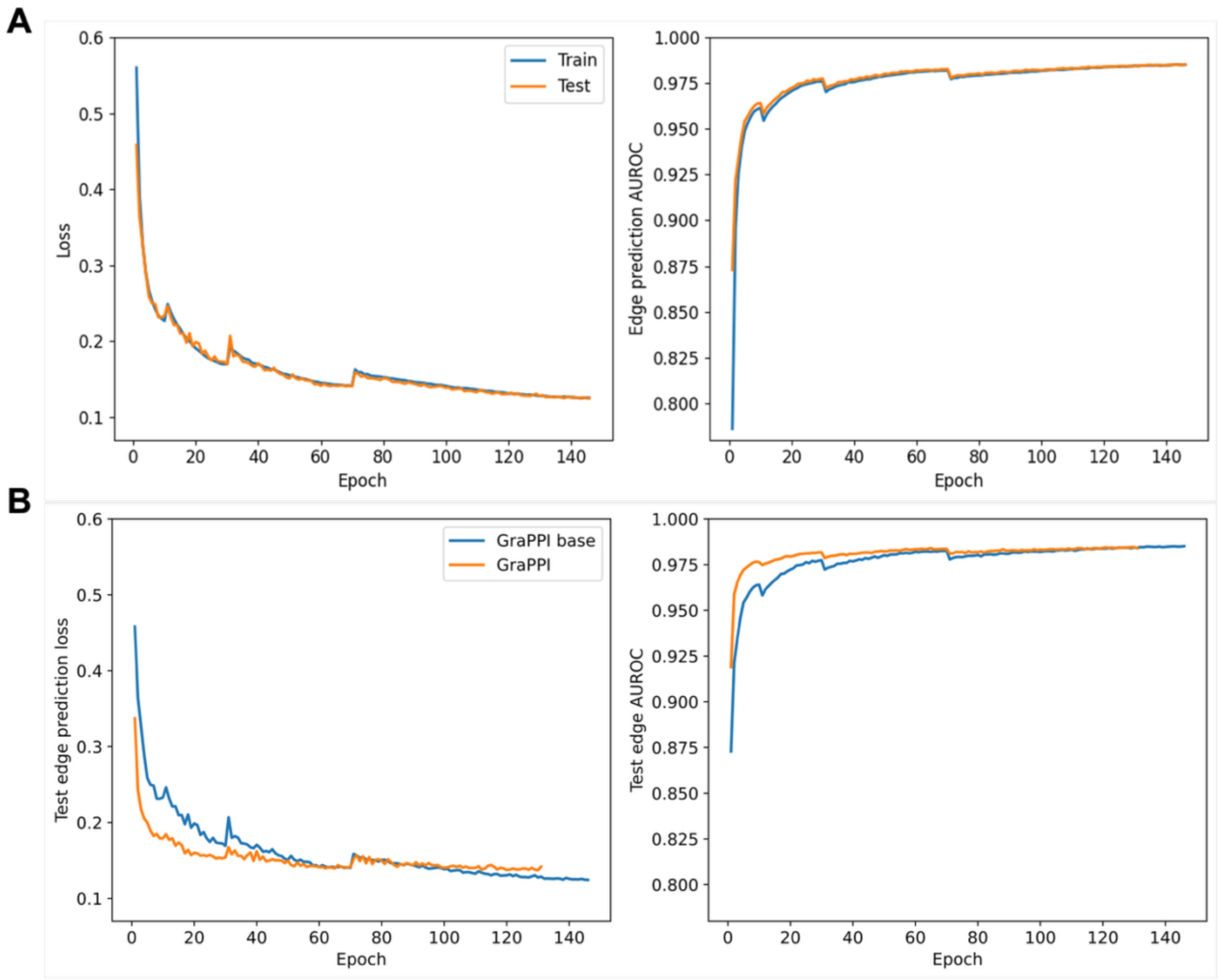


**Figure S2. Training dynamics during GraPPI self-supervised pretraining** A) Loss and AUROC over training epochs for the SSL training and validation sets. B) Test loss and AUROC of GraPPI and GraPPI-base as a function of epoch.

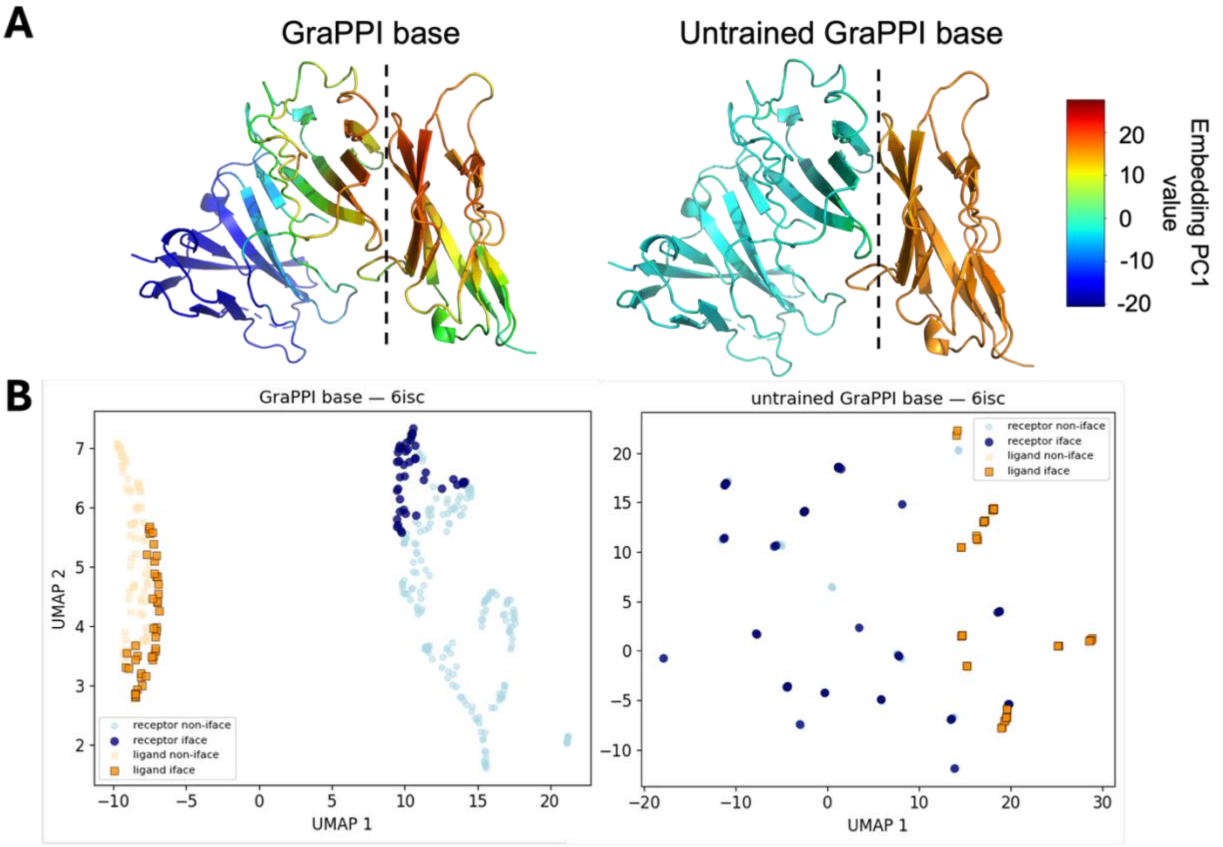


**Figure S3. Visualization of residue-level embedding representations.** (A) Residue embeddings from the immune signaling protein complex CD226-CD155 (PDB: 6ISC) were reduced to one dimension via PCA, and PC1 scores were mapped onto the protein structure to visualize the embedding patterns. The two proteins are separated by a dashed line. Results are shown for GraPPI base, untrained GraPPI base. (B) UMAP projection of residue embeddings from the CD226-CD155 complex, illustrating their organization in latent space. The two proteins are colored blue (CD226) and orange (CD155), with interface residues shown in a darker shade and non-interface residues in a lighter shade.

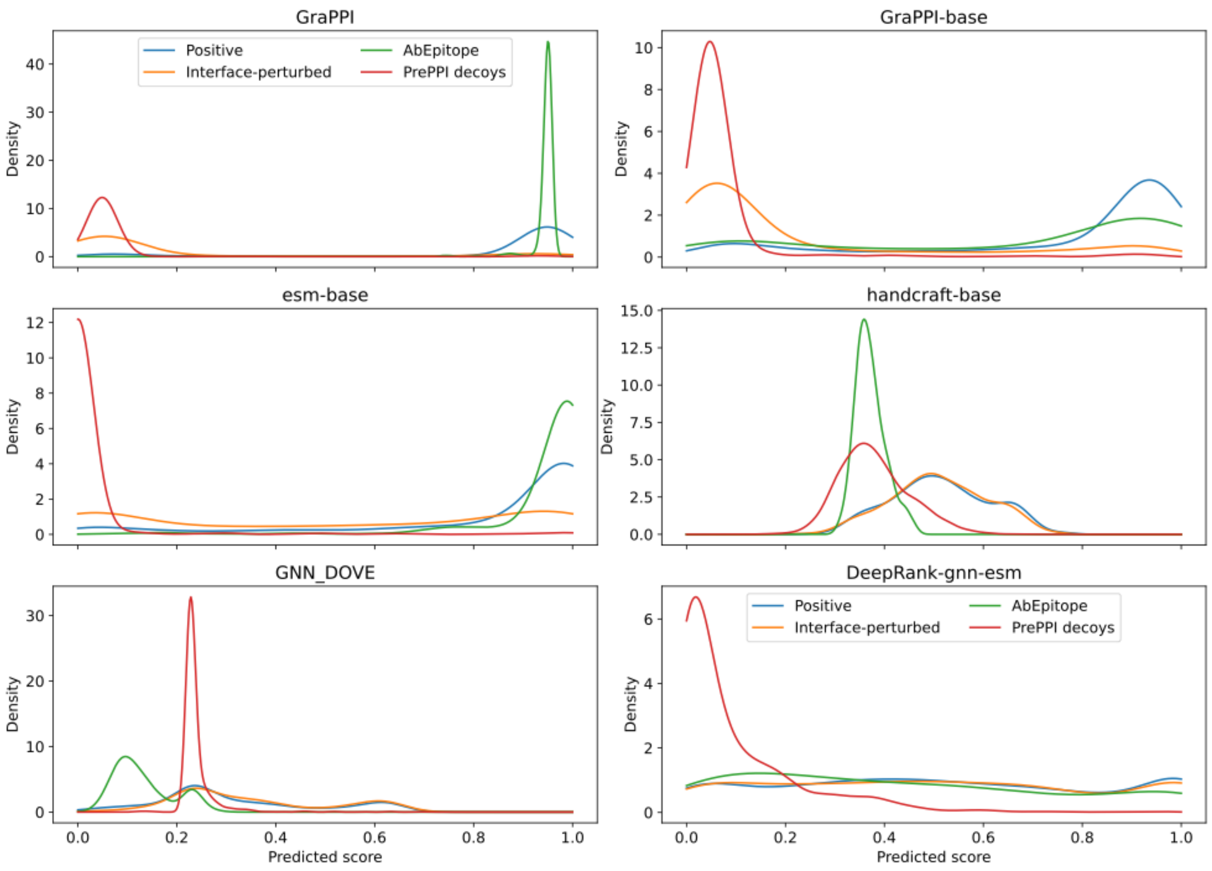


**Figure S4. Distribution of binding-mode classification scores.** Each panel shows the predicted score for a given model (GraPPI, GraPPI-base, ESM-base, Handcraft-base, GNN_DOVE, and DeepRank-gnn-esm) across four subsets: the positive, AbEpitop, interface-perturbed, and PrePPI decoy sets.

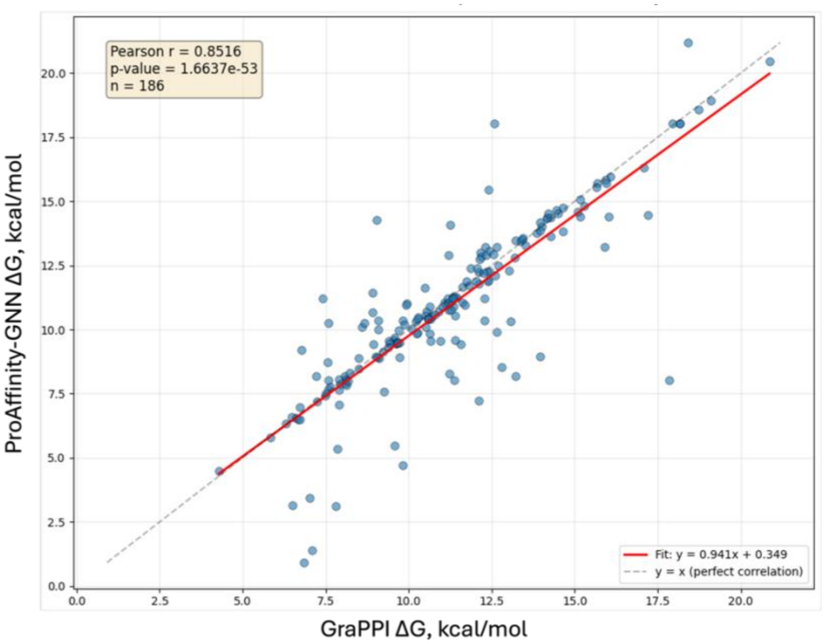


**Figure S5. Correlation between GraPPI- and ProAffinity-GNN-curated affinities for discrepant complexes.** Pearson correlation ($R_P$) between experimentally measured binding affinities as curated in the GraPPI dataset versus the ProAffinity-GNN dataset, for protein complexes where the two sources report different values.
